# Safety and Lack of Negative Effects of Wearable Augmented-Reality Social Communication Aid for Children and Adults with Autism

**DOI:** 10.1101/164335

**Authors:** Ned T. Sahin, Neha U. Keshav, Joseph P. Salisbury, Arshya Vahabzadeh

**Author notes:** Corresponding Author. Ned T. Sahin, PhD, Brain Power, 1 Broadway 14th Fl, Cambridge, MA 02142, USA.

## Abstract

There is growing interest in the use of augmented reality (AR) to assist children and adults with autism spectrum disorders (ASD); however, little investigation has been conducted into the safety of devices for AR such as smartglasses. The objective of this report was to assess the safety and negative effects of the *Brain Power Autism System* (BPAS), a novel AR smartglasses-based social communication aid for people with ASD. A sequential series of 18 children and adults aged 4.4 to 21.5 years (mean 12.2 years) with clinically diagnosed ASD of varying severity used the BPAS. Users and caregivers were interviewed about perceived negative effects and design concerns. Most users were able to wear and use the BPAS (n=16/18, 89%), with most of them reporting no negative effects (n=14/16, 87.5%). Two users reported temporary negative effects: eye strain, dizziness, and nasal discomfort. Caregivers observed no negative effects in users (n=16/16, 100%). Most users (77.8%) and caregivers (88.9%) had no design concerns. This report found no major negative effects in using an AR smartglasses-based social communication aid across a wide age and severity range of people with ASD. Further research is needed to explore longer-term effects of using AR smartglasses in this population.

## Introduction

Autism Spectrum Disorder (ASD) is a neurodevelopmental disorder affecting 1 in 68 U.S. children (1), and is characterized by social communication impairment and the presence of a restricted and/or repetitive range of interests and behaviors (2). The rising prevalence of ASD has increased the demand for educational and behavioral services, often exhausting these limited resources (3, 4). There has been considerable interest in the development and study of technology-aided approaches for the social, cognitive, and behavioral challenges related to ASD (5-7). Technology-aided approaches may be especially suitable for people with ASD given that some of these individuals may show a natural propensity to utilize digital tools (8), display a fondness for electronic media (9), express a preference for standardized and predictable interactions (8), enjoy game-like elements (10) and/or favor computer-generated speech (11). However, technology may also have negative effects in some people with ASD. Individuals may develop problematic video game use (12), and can become agitated or disruptive when attempting to disengage from video games (13). Anecdotally, many caregivers describe meltdowns and other episodes of behavioral dysregulation in children with ASD when attempting to stop them playing on smartphone and/or tablets (14).

Evidence suggests that a broad range of technology-aided interventions, such as those using computer programs and virtual reality (VR), may be effective in people with ASD (5). Technology-based interventions have been found to be beneficial for improving a wide range of skills and behaviors including aiding social and emotional skills (15-17), communication skills (16), academic skills (18), employment skills (6), and challenging behaviors (15).

There is particular interest in interventions that help users learn skills while continuing to interact with the people and environment around them. Learning social-emotional skills in real life settings (such as in social skills groups), may increase the chance these skills generalize to the challenges of daily life (19). Augmented reality (AR) is a technology that holds considerable promise in this regard, as it allows for users to see and interact with the real world around them, while virtual objects and audio guidance are provided through a visual overlay and audio speakers (**Figure 1A**). In contrast, current VR headsets place users and their senses into an entirely virtual world, while simultaneously removing their ability to see and sense real-world stimuli, hazards, and social situations around them (**Figure 1C**). In contrast to VR headsets, AR allows users to see their real-world environment, allowing for them to more readily navigate an environmental hazard, or to socially engage with another person. Nonetheless, AR incorporates many of the features of VR that are thought to make VR technology well suited to the creation of learning tools for people with ASD (20), including being a primarily visual and auditory experience, being able to individualize the experience, and promoting generalization and decreasing rigidity through subtle gradual modifications in the experience (20).

**Figure 1.**
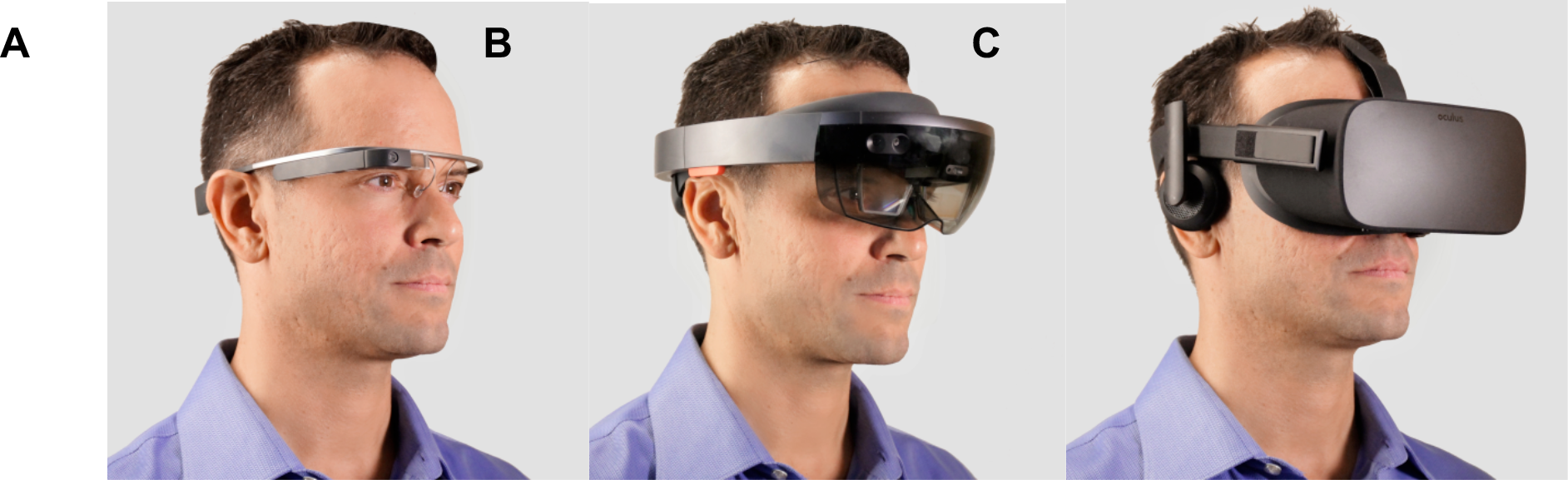
Head-worn Computers or Displays. (**A**) Glass Explorer Edition (originally known as Google Glass): augmented reality (AR) smartglasses with fully stand-alone onboard computer (weight 42 grams). (**B**) Microsoft Hololens: AR headset with fully stand-alone onboard computer and depth camera (weight 579 grams). (**C**) Oculus Rift: virtual reality (VR) headset display, which must be tethered continuously to a powerful computer to drive it (weight 470 grams). VR headsets and some AR devices are large, heavy, and block the social world considerably.

AR experiences can also be easily modified and personalized for each individual, an important consideration given that many people with ASD exhibit intense interest in a restricted range of topics, and may experience extreme distress if changes to their routine/environment occur (2). AR experiences are also not restricted solely to real-world limitations on time, space, and resources. For instance, users may have the opportunity to interact with objects or experiences from historical or fantasy worlds, or a simplified and cartoon-like interaction where the sensory and perceptual experiences may be reduced in complexity and/or magnitude.

Most ASD-related research into AR has focused on the use of smartphone- and/or tablet-based apps. While research has been limited, AR apps on smartphones/tablets have been shown to improve selective and sustained attention (21), attention to social cues (22), the ability to understand emotions and facial expressions in storybook characters (22), and navigating the physical world when attempting to find employment opportunities (23). However, smartphone based AR may carry with it a risk of negative effects, including grip and postural strain, minor falls, and falls leading to major trauma and blindness (24, 25).

While AR has been investigated as an educational medium in ASD children for at least a decade (26), minimal research has been conducted into the safety of head-mounted AR in ASD populations. This has potential implications as head-mounted AR, in particular smartglasses, may offer advantages compared to smartphone- or tablet-based AR, and may be the optimal future platform for AR (27). Generalized use of AR smartglasses may still be in its infancy, but use of such devices will be fueled by their ability to improve social interactions and relationships, make life more efficient, and provide enjoyment and fun to the user (28). AR smartglasses may also be beneficial tools for clinical research. AR smartglasses contain a wide array of sensors. These are intended to allow basic features such as gesture-based control of the devices (to make up for the lack of keyboards and traditional input devices). However, we have shown that these sensors can also be used creatively to collect quantitative data that may help assess brain function (29). Analysis of quantitative data from sensors in smart-devices may help to advance digital phenotyping of neurobehavioral conditions (30). To our knowledge, we have published the first reports of the use of AR smartglasses in children with ASD (30-33). Even in terms of VR, about which there are generally many more reports in the literature, there are very few reports of people with ASD using modern VR headsets (34, 35).

Therefore, it would be useful to understand how children and adults with ASD would respond to AR smartglasses, particularly when the smartglasses function as an assistive device loaded with specialized assistive social and behavioral coaching software (30). Of first importance in assessing a new assistive technology is to assess a.) the safety of such an approach, and b.) any potential negative effects.

There are both human and device factors that make it conceivable that even commercially-available AR technology could elicit concerns regarding safety or negative effects, when applied as an assistive technology for this special population.

Firstly, in regard to human factors, it has been widely reported that people with ASD have a range of sensory (2, 36), motor (37), and cognitive challenges (38, 39), as well as strong negative reactions to transitions (40). More specifically, ASD is often accompanied by atypical reactivity to sensory inputs such as touch, sound, temperature, and sight (2). Altered sensory reactivity is not only prevalent but also highly heterogeneous in the ASD population. Each member of this diverse spectrum may be affected in none, one, or several senses, and as a hyper- or hypo-sensitivities, representing a complex matrix of sensory subtypes (36). It is therefore important to measure if individuals can safely use smartglasses for an extended period, and to monitor how they respond to visual, auditory, and vibrotactile cues delivered through the device.

Secondly, there may be safety concerns because ASD is often associated with altered motor movements, such as repetitive behaviors (a “core” symptom of ASD)(2) or impairments of motor coordination (37). It is thus important to assess if such motor challenges may lead to falls or injury when people with ASD utilize AR smartglasses (37).

Thirdly, people with ASD may differ in their ability to remain attentive and to focus on using smartglasses as part of social communication training, especially given the high rate of comorbidity between ASD and attention deficit hyperactivity disorder (ADHD)(41). Some individuals may find themselves becoming distracted when using AR, or in the process of becoming familiar with using AR while simultaneously navigating the real world (42).

These attentional difficulties may compound the motor coordination challenges in ASD as mentioned above, to increase the potential of AR smartglasses use to result in falls and/or trips. Additionally, over 30% of children with ASD demonstrate wandering/elopement behavior, and it would be prudent to investigate any technology that would affect their perception, attention, and ability to avoid hazards (43).

Finally, people with ASD may face major challenges in coping with transitions in activities (2, 44) and have demonstrated oppositional defiant behaviors and aggression when asked to stop playing video games (13) or using piece of technology (14). This suggests a possible risk of meltdown when an AR session is brought to an end, though it remains to be seen whether stopping use of smartglasses results in less difficulty than when stopping use of a smartphone or tablet (which may be more engrossing or cognitively demanding).

Instruction manuals for AR smartglasses are an additional indication that there may be device-related factors that result in risks. For instance, the Microsoft HoloLens manual identifies potential side effects as being nausea, motion sickness, dizziness, disorientation, headache, fatigue, eye strain, dry eyes, and seizures (45), although their occurrence among users with ASD has not been studied. Little study has investigated how these new AR devices may impact the perceptual abilities of regular users, raising concerns that some individuals may become distracted, have altered reaction times, misjudge hazards in the real-world, and/or experience altered distance and speed perception (42).

AR may share a subset of the risks of VR, and VR research has reported potential side effects that include eye strain, headache, and disorientation during use of a VR headset (46). However, there have been continuing advances in VR technology, and a recent study noted that people with ASD experienced relatively few negative effects when using a VR headset of the modern generation (35).

Assessing negative effects in people with ASD is not a simple undertaking given that these individuals have challenges in communicating their experiences. It is therefore important to explicitly ask for their feedback, and also seek feedback from their caregivers in order to have a more comprehensive method of detecting any negative effects.

## Aims of Research

Given the potential for AR smartglasses to be used in people with ASD, and yet the uncertainty as to whether this technology would be safe in this population, we studied a specific AR smartglasses technology in 18 children and adults with ASD. The system used in this study was the *Brain Power Autism System* (BPAS)(30).

## Brain Power Autism System (BPAS)

BPAS is a social communication aid that consists of AR smartglasses with apps that allow children and adults with ASD to coach themselves on important socio-emotional and cognitive skills (30-32). Users learn through gamified interactions and a combination of intrinsic and extrinsic rewards for successfully completing tasks. In certain situations, such as coaching of appropriate face-directed gaze and emotion recognition, the BPAS is designed to be used while the user is interacting with another person. The system was designed using an iterative process where continuous feedback from people with ASD, clinicians, neuroscientists, educators, caregivers, design specialists, and engineers, helped to develop the system that was used in this report. Additionally, the facial affective analytics component of the BPAS was developed in partnership with Affectiva, an emotion artificial intelligence company. The work was also made possible by Google, Inc., now known as Alphabet, Inc., who provided substantial hardware as well as guidance in engineering. Engineering guidance, including as to how best to develop apps that would be accessible to a diverse set of users, was provided in part in the context of the Glass Enterprise Partnership Program. BPAS is designed to be accessible to people with ASD, and to minimize potential negative effects. A number of elements were used to achieve this, including but not limited to the use of calming tones, use of emotional artificial intelligence, minimization of audio and visual sensory load, graduated transitions between learning segments, and modification of the functionality of the tactile input surfaces of the smartglasses. In this study the focus was on understanding the safety and potential negative effects that may be experienced by children and adults with ASD as they used AR smartglasses delivering cognitive and social self-coaching apps.

## Methods

The methods and procedures of this study were approved by Asentral, Inc., Institutional Review Board, an affiliate of the Commonwealth of Massachusetts Department of Public Health.The study was performed in accordance with relevant guidelines and regulations.

### User Recruitment

A sequential sample of 18 children and adults with ASD were recruited from a database of individuals who completed a web-based signup form expressing interest in our study (mean age 12.2 years, range: 4.4 – 21.5 years; **Table 1**). Users included males and females, both verbal and non-verbal, and represented a wide range of ASD severity levels. Users had Social Communication Questionnaire scores from 6 to 28, with an average of 18.8. The Social Communication Questionnaire is a validated way to obtain diagnostic and screening information about ASD (47, 48). Written informed consent was obtained from the legal guardians of children and from cognitively able adults. Participants between 7 and 17 years-old additionally provided written assent, when possible. In this report, every user was accompanied by a caregiver, and users and caregivers could exit the session at any time and for any reason.

**Table 1.** Demographics of Study Participants.

| Demographics |  |  |
| --- | --- | --- |
| Number of users | 18 |  |
| Age (mean +/- SD) | 12.2 +/- 5.2 | Range = 4.4 years – 21.5 years |
| Participant gender | Male: 16 (88.9%) | Female: 2 (11.1%) |
| Verbal or nonverbal | Verbal: 16 (88.9%) | Nonverbal: 2 (11.1%) |
| Social Communication Questionnaire (SCQ) Score (mean +/- SD) | 18.8 +/- 6.75 | Range = 6 – 28 |

### Exclusions

Individuals who had expressed interest via the website signup but who had a known history of epilepsy or seizure disorder were not enrolled in this study. Users who had an uncontrolled or severe medical or mental health condition that would make participation in the study very difficult were also not enrolled.

### Data Collection Procedure

Users and caregivers had the opportunity to wear the BPAS, while they were video and audio recorded. In this report, the base smartglasses technology underlying the BPAS was Glass Enterprise Edition (originally known as Google Glass) (**Figure 2**). Users who could physically wear the smartglasses for at least one minute were allowed to proceed to trying the different BPAS social and cognitive coaching apps. They used the apps over a period of 1–1.5 hours. The level of variability in the session length required to use the range of apps was reflective of the considerable range of ASD severity in the user group. Users interacted with their caregivers while they practiced with the apps, and were required to frequently take off the smartglasses and then put them back on (**Figure 3**). Following the experience with the system, structured interviews were conducted with users and their caregivers. In the structured interviews, users and caregivers were asked to identify any perceived negative effects of using the system, and were allowed to raise concerns or give comments about the design of the smartglasses hardware as well as the apps.

**Figure 2.**
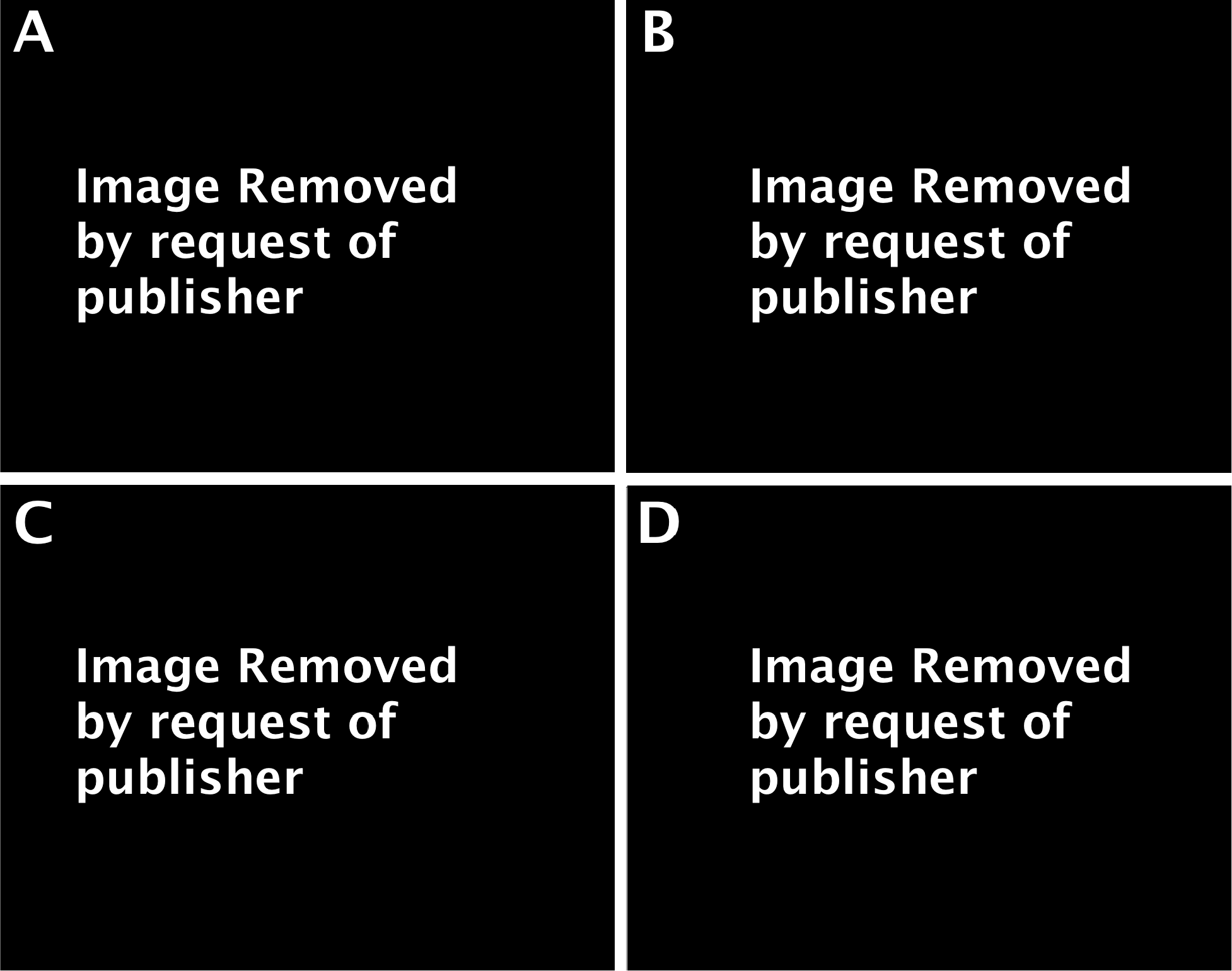
Smartglasses Platform in Use. Four representative trial participants wearing the Brain Power Autism System (BPAS). This version of the BPAS used the Glass Explorer Edition device (originally known as Google Glass). All pictures are used with user / caregiver consent.

**Figure 3.**
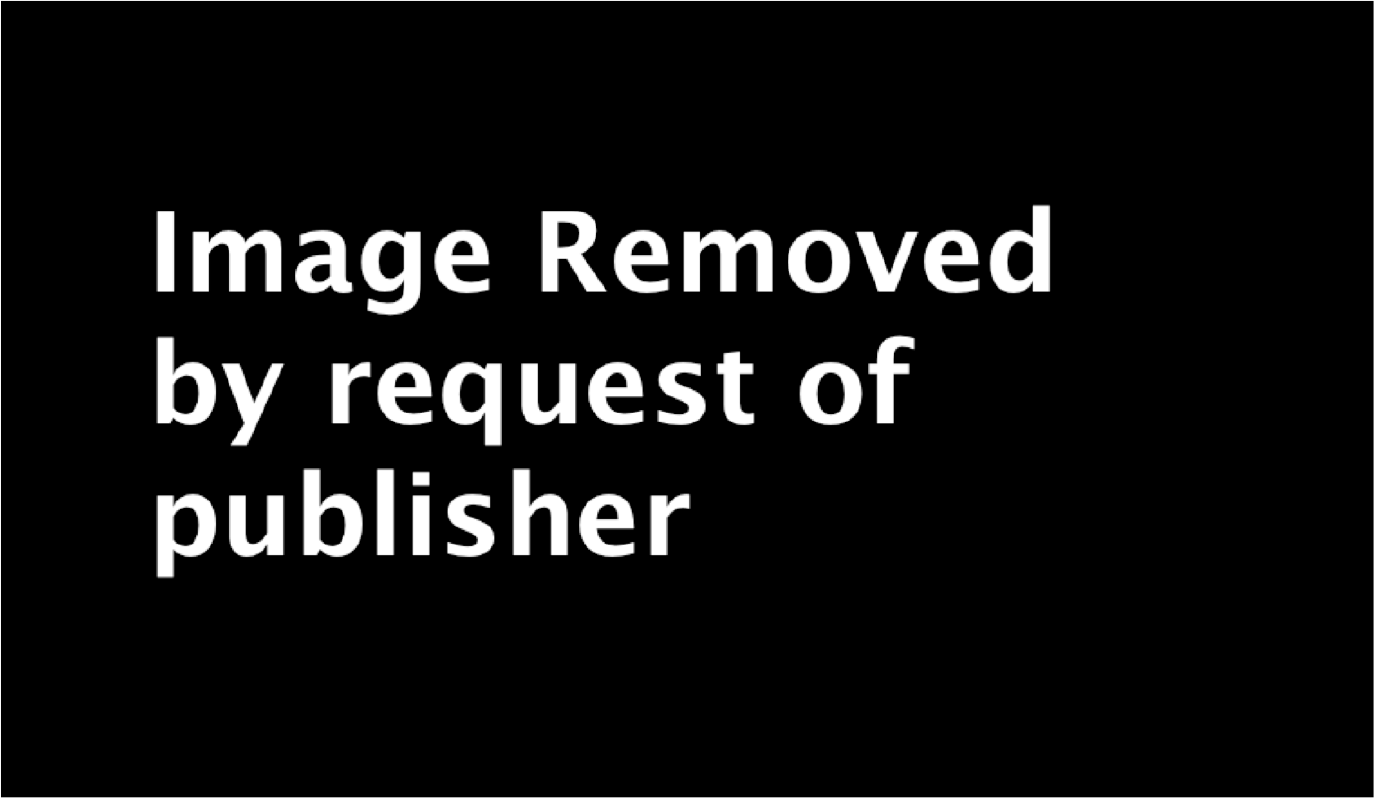
BPAS User-Caregiver Setup. In each session, the participant and caregiver sit facing one another, promoting ‘heads-up’ social interaction while trialing BPAS apps. All pictures are used with user / caregiver consent.

## Results

Sixteen of the 18 users (89%) tolerated wearing BPAS smartglasses for at least one minute. The two users who did not tolerate this initial testing did not use BPAS apps. While both of these two users and their caregivers did not report any adverse effects, the users did not express an interest in wearing the BPAS or continuing the testing session. It was noted that both users were non-verbal, and were relatively young, aged 5.5 and 5.8 years. Of the remaining users, 14 out of 16 users (87.5%), and 16 out of 16 caregivers (100%) reported no minor negative effects, and 100% of caregivers and users reported no major negative effects (**Table 2**)(**Figure 4**).

**Table 2.**
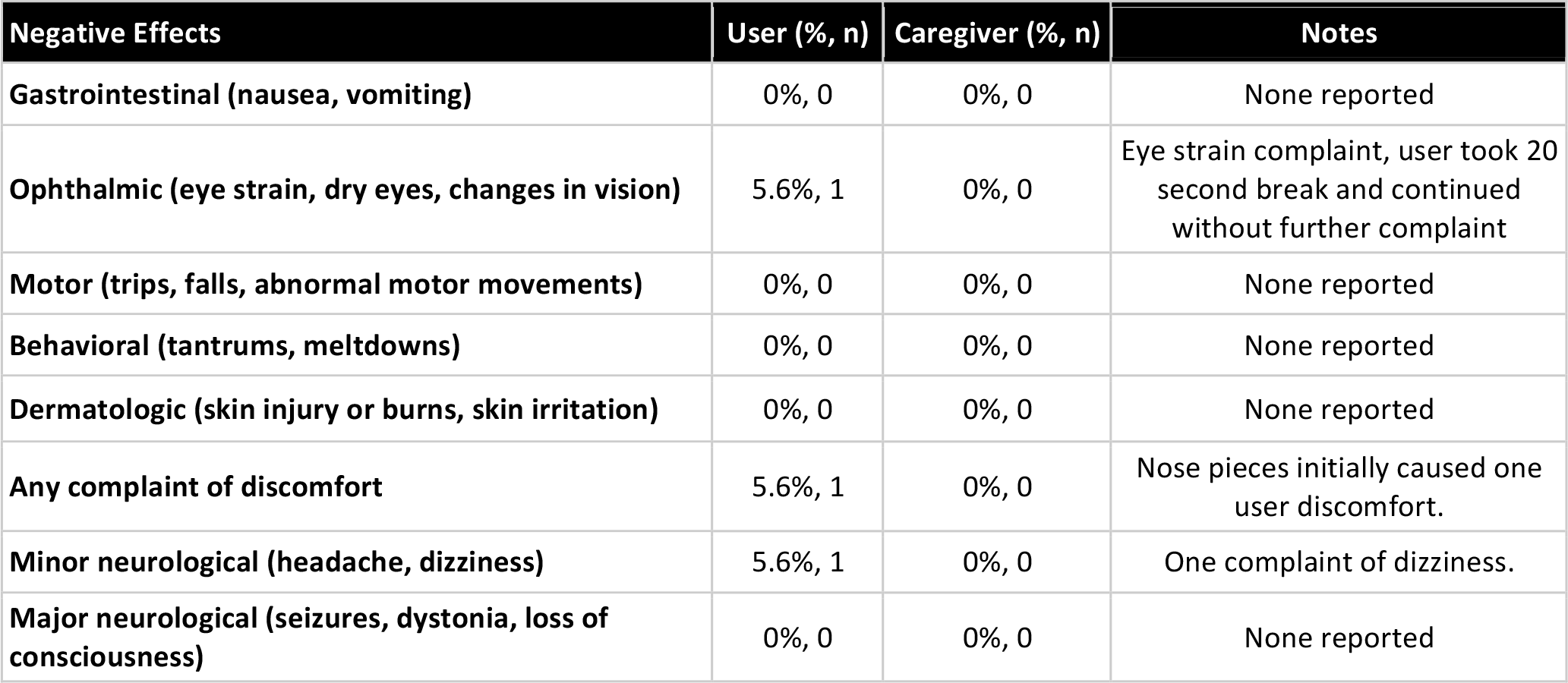
Negative Effects.

| Negative Effects | User (% , n) | Caregiver (% , n) | Notes |
| --- | --- | --- | --- |
| Gastrointestinal (nausea, vomiting) | 0%, 0 | 0%, 0 | None reported |
| Ophthalmic (eye strain, dry eyes, changes in vision) | 5.6%, 1 | 0%, 0 | Eye strain complaint, user took 20 second break and continued without further complaint |
| Motor (trips, falls, abnormal motor movements) | 0%, 0 | 0%, 0 | None reported |
| Behavioral (tantrums, meltdowns) | 0%, 0 | 0%, 0 | None reported |
| Dermatologic (skin injury or burns, skin irritation) | 0%, 0 | 0%, 0 | None reported |
| Any complaint of discomfort | 5.6%, 1 | 0%, 0 | Nose pieces initially caused one user discomfort. |
| Minor neurological (headache, dizziness) | 5.6%, 1 | 0%, 0 | One complaint of dizziness. |
| Major neurological (seizures, dystonia, loss of consciousness) | 0%, 0 | 0%, 0 | None reported |

**Figure 4.**
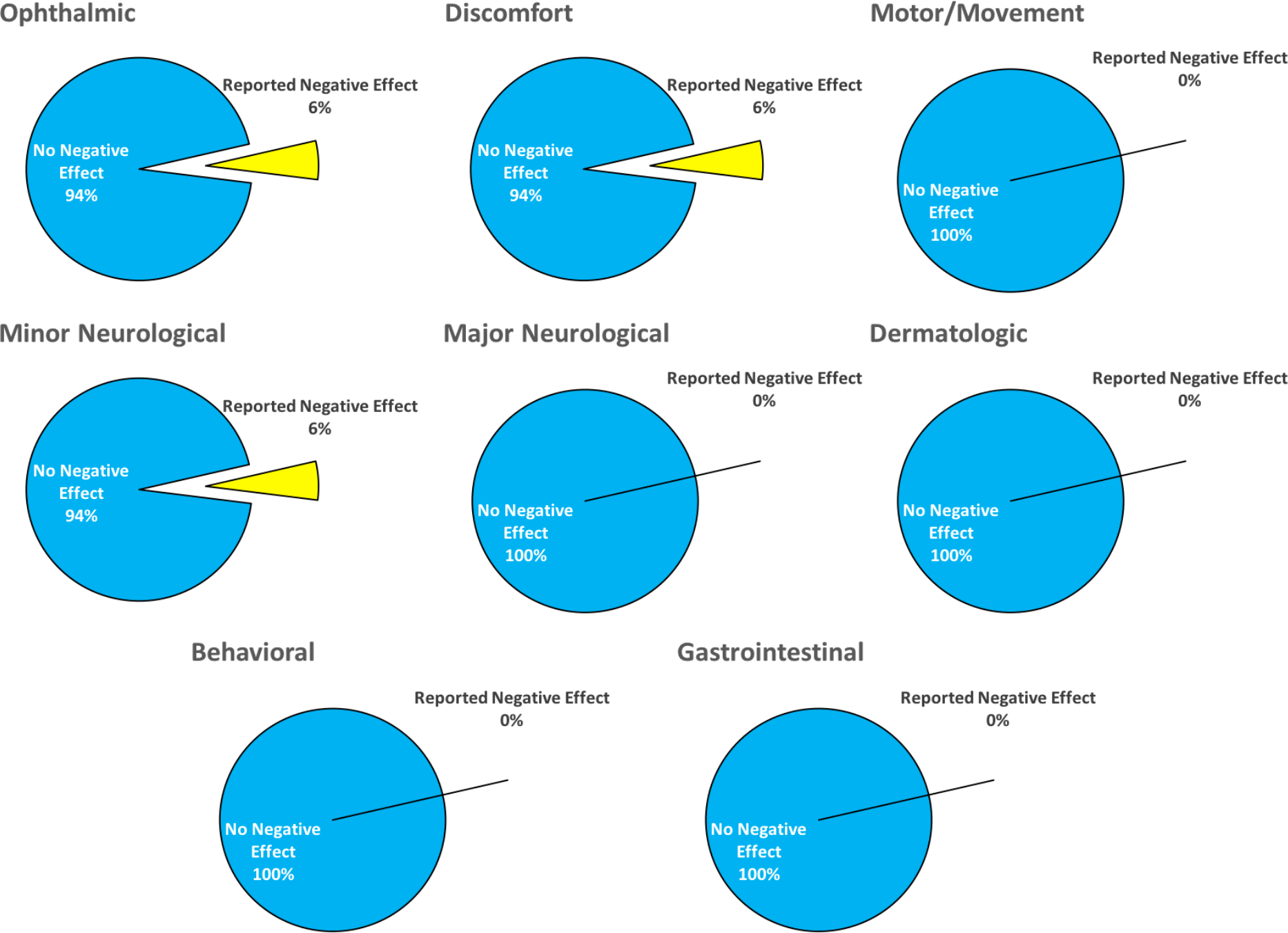
User-Reported Negative Effects by Category. The three negative effects that were reported (by two participants) include one episode of eye strain (Ophthalmic), one episode of nasal bridge discomfort (Discomfort), and one episode of dizziness (Minor Neurological).

The three instances of negative effects were reported by 2 users. The effects were all mild in nature, transitory in duration, and did not result in the session being stopped. The reported negative effects were one case of dizziness, one case of eye strain, and one instance of initial nasal bridge discomfort. The caregiver of the user experiencing dizziness later explained that the effect may have been related to the user not wearing his prescription glasses, and that he had previously experienced similar dizziness when he had tried a modern VR headset. This same user also experienced initial discomfort with the nose pads, but resolved the discomfort with adjustment of the placement of the smartglasses. The user who had complained of eye strain resolved the issue with a 20-second break in testing.

The majority of users and caregivers did not have any design concerns about the system (77.8% and 88.9% respectively) (**Table 3**). The only design concern highlighted by users and caregivers was that the smartglasses became warm to the touch during use, although this did not result in any negative effects.

**Table 3.**
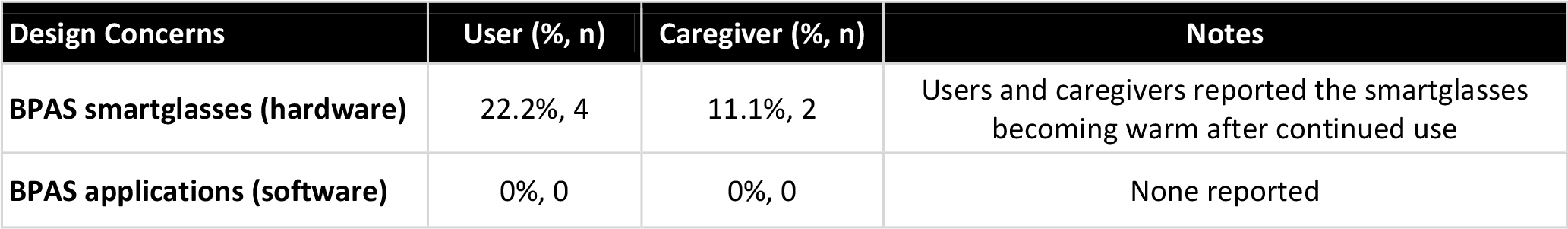
Design concerns. Design concerns reported by users and by caregivers, including concerns raised spontaneously during or following testing session, as well as those mentioned in response to direct questions about design during structured interviews following testing sessions.

| Design Concerns | User (% , n) | Caregiver (% , n) | Notes |
| --- | --- | --- | --- |
| BPAS smartglasses (hardware) | 22.2%, 4 | 11.1%, 2 | Users and caregivers reported the smartglasses becoming warm after continued use |
| BPAS applications (software) | 0%, 0 | 0%, 0 | None reported |

## Discussion

The safety and effects of AR smartglasses in children and adults with ASD is an important but poorly researched area, especially given the potential benefits of this technology. People with ASD who use AR smartglasses could potentially experience negative effects due to a range of known device-related factors and ASD-related human factors. ASD-related human factors include challenges in sensory, motor, attentional, and transition-related process. Device-related factors, as per manufacturer warnings about side effects, include dizziness, headache, and seizures.

This paper explored the use of the BPAS, a novel technological system that uses AR smartglasses to deliver social and cognitive coaching to children and adults with ASD. The focus was on exploring safety and negative effects of using this technology across a broad age and severity range of children and adults with clinically diagnosed ASD. The practicalities of conducting this research involved circumstances that the authors believe could have made the experience more difficult for users than would have been the case had they tested/used the AR smartglasses in a more naturalistic home setting. During the day of testing, users and caregivers were exposed to novel surroundings by attending the research center, asked to undertake a number of environmental transitions prior to the testing, and users had the additional sensory load of being video and audio recorded while using the BPAS.

In this context our results are encouraging, and suggest that the majority of people with ASD can use AR smartglasses without reporting any negative effects. Of the 16 users who managed to wear and use the BPAS (n=16/18), caregiver and user reports found no negative effects in 14 (n=14/16, 87.%). In the 2 individuals who reported negative effects, there were 3 reported instances of negative effects: one case of dizziness, one case of eye strain, and one instance of initial nasal bridge discomfort. These negative effects were mild in nature, temporary, and did not lead to the user or caregiver stopping the session. It was interesting to see that in these cases the caregiver had not reported these negative effects, therefore justifying the explicit interviewing of people with ASD in order to understand their experience of this technology. As we previously noted, two individuals were not able to, or did not show interest in wearing/using the BPAS. These 2 individuals were both noted to be non-verbal and relatively young, and were not reported to have had any negative effects. This suggests that a small group of people with ASD may struggle to utilize current AR smartglasses.

The relative lack of negative effects in this AR paradigm is an important finding across such a wide age and severity range of people with ASD, and indirectly supports recent research demonstrating minimal negative effects when modern VR headsets were used by people with ASD (35). It was reassuring to see that no major negative effects were reported, and additionally no behavioral problems such as tantrums or meltdowns occurred when users were asked to stop using the smartglasses, especially given earlier outlined concerns regarding the potential for distress around transitions involving technology.

There were no design concerns by the majority of caregivers and users. Design concerns were raised by 2 caregivers and 4 users who noticed a feeling of warmth from the external side of the hardware after extended use. However, this did not result in any reported negative effects. User acceptance of design is an important part of any assistive technology experience, so it was useful to know that users and caregivers had few concerns about the design and use of the BPAS.

There are a number of limitations to our findings. The results may not generalize to systems based on other types of AR smartglasses, or that include other software apps. There remains a critical need to conduct further research to understand the feasibility and safety associated with new emerging technologies, especially those that may be used in vulnerable populations such as ASD. The use of AR smartglasses may have considerable potential as an augmentative technology in helping people with ASD, particularly when they are shown to be usable and safe in the ASD population, and when supported by robust evidence of efficacy. Finally, while this report does not identify any short term adverse events, as with any technology, further research is warranted to explore the positive and negative effects of longer term use.

## Author Contributions

NTS is the inventor of the Brain Power Autism System (BPAS). NTS was the primary author of this paper with additional review and comments by NUK, JPS, and AV. NTS, NUK, JPS, and AV helped to design and conduct the study.

## REFERENCES

1. Developmental DMNSY, Investigators P. Prevalence of autism spectrum disorder among children aged 8 years-autism and developmental disabilities monitoring network, 11 sites, United States, 2010. Morbidity and mortality weekly report Surveillance summaries (Washington, DC: 2002). 2014;63(2):1.

2. Association AP. Diagnostic and Statistical Manual of Mental Disorders (DSM-5®): American Psychiatric Pub; 2013.

3. Kogan MD, Strickland BB, Blumberg SJ, Singh GK, Perrin JM, van Dyck PC. A national profile of the health care experiences and family impact of autism spectrum disorder among children in the United States, 2005–2006. Pediatrics. 2008;122(6):e1149–e58.

4. Boyle CA, Boulet S, Schieve LA, Cohen RA, Blumberg SJ, Yeargin-Allsopp M, et al. Trends in the prevalence of developmental disabilities in US children, 1997–2008. Pediatrics. 2011;127(6):1034–42.

5. Grynszpan O, Weiss PL, Perez-Diaz F, Gal E. Innovative technology-based interventions for autism spectrum disorders: a meta-analysis. Autism. 2014;18(4):346–61.

6. Walsh E, Holloway J, McCoy A, Lydon H. Technology-Aided Interventions for Employment Skills in Adults with Autism Spectrum Disorder: A Systematic Review. Review Journal of Autism and Developmental Disorders. 2017:1–14.

7. Didehbani N, Allen T, Kandalaft M, Krawczyk D, Chapman S. Virtual reality social cognition training for children with high functioning autism. Computers in Human Behavior. 2016;62:703–11.

8. Golan O, Baron-Cohen S. Systemizing empathy: Teaching adults with Asperger syndrome or high-functioning autism to recognize complex emotions using interactive multimedia. Development and psychopathology. 2006;18(02):591–617.

9. Shane HC, Albert PD. Electronic screen media for persons with autism spectrum disorders: Results of a survey. Journal of autism and developmental disorders. 2008;38(8):1499–508.

10. Mayes SD, Calhoun SL. Analysis of WISC-III, Stanford-Binet: IV, and academic achievement test scores in children with autism. Journal of autism and developmental disorders. 2003;33(3):329–41.

11. Schlosser RW, Blischak DM. Is there a role for speech output in interventions for persons with autism? A review. Focus on Autism and Other Developmental Disabilities. 2001;16(3):170–8.

12. Mazurek MO, Engelhardt CR. Video game use in boys with autism spectrum disorder, ADHD, or typical development. Pediatrics. 2013;132(2):260–6.

13. Mazurek MO, Engelhardt CR. Video game use and problem behaviors in boys with autism spectrum disorders. Research in Autism Spectrum Disorders. 2013;7(2):316–24.

14. Samuel A. Personal Technology and the Autistic Child: What One Family Has Learned. 2017.

15. Ramdoss S, Lang R, Fragale C, Britt C, O’Reilly M, Sigafoos J, et al. Use of computer-based interventions to promote daily living skills in individuals with intellectual disabilities: A systematic review. Journal of Developmental and Physical Disabilities. 2012;24(2):197–215.

16. Ramdoss S, Lang R, Mulloy A, Franco J, O’Reilly M, Didden R, et al. Use of computer-based interventions to teach communication skills to children with autism spectrum disorders: A systematic review. Journal of Behavioral Education. 2011;20(1):55–76.

17. Ramdoss S, Machalicek W, Rispoli M, Mulloy A, Lang R, O’Reilly M. Computer-based interventions to improve social and emotional skills in individuals with autism spectrum disorders: A systematic review. Developmental neurorehabilitation. 2012;15(2):119–35.

18. Pennington RC. Computer-assisted instruction for teaching academic skills to students with autism spectrum disorders: A review of literature. Focus on Autism and Other Developmental Disabilities. 2010;25(4):239–48.

19. Reichow B, Steiner AM, Volkmar F. Cochrane review: social skills groups for people aged 6 to 21 with autism spectrum disorders (ASD). Evidence-Based Child Health: A Cochrane Review Journal. 2013;8(2):266–315.

20. Strickland D. Virtual reality for the treatment of autism. Studies in health technology and informatics. 1997:81–6.

21. Escobedo L, Tentori M, Quintana E, Favela J, Garcia-Rosas D. Using augmented reality to help children with autism stay focused. IEEE Pervasive Computing. 2014;13(1):38–46.

22. Chen C-H, Lee I-J, Lin L-Y. Augmented reality-based video-modeling storybook of nonverbal facial cues for children with autism spectrum disorder to improve their perceptions and judgments of facial expressions and emotions. Computers in Human Behavior. 2016;55:477–85.

23. McMahon D, Cihak DF, Wright R. Augmented reality as a navigation tool to employment opportunities for postsecondary education students with intellectual disabilities and autism. Journal of Research on Technology in Education. 2015;47(3):157–72.

24. Radu I, Guzdial K, Avram S, editors. An Observational Coding Scheme for Detecting Children’s Usability Problems in Augmented Reality. Proceedings of the 2017 Conference on Interaction Design and Children; 2017: ACM.

25. Franchina M, Sinkar S, Ham B, Lam GC. A blinding eye injury caused by chasing Pokemon. Med J Aust. 2017;206(9):384.

26. Richard E, Billaudeau V, Richard P, Gaudin G, editors. Augmented reality for rehabilitation of cognitive disabled children: A preliminary study. Virtual Rehabilitation, 2007; 2007: IEEE.

27. Monkman H, Kushniruk AW, editors. A see through future: augmented reality and health information systems. ITCH; 2015.

28. Rauschnabel PA, Brem A, Ro Y. Augmented reality smart glasses: definition, conceptual insights, and managerial importance. Unpublished Working Paper, The University of Michigan-Dearborn, College of Business. 2015.

29. Salisbury JP, Keshav NU, Sossong AD, Sahin NT. Standing balance assessment using a head-mounted wearable device. bioRxiv. 2017.

30. Liu R, Salisbury JP, Vahabzadeh A, Sahin NT. Feasibility of an Autism-Focused Augmented Reality Smartglasses System for Social Communication and Behavioral Coaching. Frontiers in Pediatrics. 2017;5(145).

31. Vahabzadeh A, Keshav NU, Salisbury JP, Sahin NT. Preliminary Report on the Impact of Smartglasses-based Behavioral and Social Communication Aid on Hyperactivity in Children and Adults with Autism. bioRxiv. 2017.

32. Keshav NU, Salisbury JP, Vahabzadeh A, Sahin NT. But will they even wear it? Exploring the tolerability of social communication coaching smartglasses in children and adults with autism. bioRxiv. 2017.

33. Sahin NT, Keshav NU, Salisbury JP, Vahabzadeh A. Cool Enough for School: Second Version of Google Glass Rated by Children Facing Challenges to Social Integration as Desirable to Wear at School. bioRxiv. 2017.

34. Newbutt N, Sung C, Kuo HJ, Leahy MJ. The acceptance, challenges, and future applications of wearable technology and virtual reality to support people with autism spectrum disorders. Recent Advances in Technologies for Inclusive Well-Being: Springer; 2017. p. 221–41.

35. Newbutt N, Sung C, Kuo H-J, Leahy MJ, Lin C-C, Tong B. Brief report: a pilot study of the use of a virtual reality headset in autism populations. Journal of autism and developmental disorders. 2016;46(9):3166–76.

36. Ausderau KK, Sideris J, Little LM, Furlong M, Bulluck JC, Baranek GT. Sensory subtypes and associated outcomes in children with autism spectrum disorders. Autism Research. 2016;9(12):1316–27.

37. Fournier KA, Hass CJ, Naik SK, Lodha N, Cauraugh JH. Motor coordination in autism spectrum disorders: a synthesis and meta-analysis. Journal of autism and developmental disorders. 2010;40(10):1227–40.

38. Howlin P, Savage S, Moss P, Tempier A, Rutter M. Cognitive and language skills in adults with autism: a 40-year follow-up. Journal of Child Psychology and Psychiatry. 2014;55(1):49–58.

39. Landry O, Al-Taie S. A meta-analysis of the Wisconsin Card Sort Task in autism. Journal of autism and developmental disorders. 2016;46(4):1220–35.

40. Sevin JA, Rieske RD, Matson JL. A Review of Behavioral Strategies and Support Considerations for Assisting Persons with Difficulties Transitioning from Activity to Activity. Review Journal of Autism and Developmental Disorders. 2015;2(4):329–42.

41. Ronald A, Simonoff E, Kuntsi J, Asherson P, Plomin R. Evidence for overlapping genetic influences on autistic and ADHD behaviours in a community twin sample. Journal of Child psychology and Psychiatry. 2008;49(5):535–42.

42. Sabelman EE, Lam R. The real-life dangers of augmented reality. IEEE Spectrum. 2015;52(7):48–53.

43. Rice CE, Zablotsky B, Avila RM, Colpe LJ, Schieve LA, Pringle B, et al. Reported wandering behavior among children with autism spectrum disorder and/or intellectual disability. The Journal of pediatrics. 2016;174:232–9. e2.

44. Dettmer S, Simpson RL, Myles BS, Ganz JB. The use of visual supports to facilitate transitions of students with autism. Focus on Autism and Other Developmental Disabilities. 2000;15(3):163–9.

45. Microsoft. Health & Safety https://www.microsoft.com/en-us/hololens/legal/health-and-safety-information: Microsoft; 2017 [

46. LaViola JR JJ. A discussion of cybersickness in virtual environments. ACM SIGCHI Bulletin. 2000;32(1):47–56.

47. Rutter M, Bailey A, Lord C. The social communication questionnaire: Manual: Western Psychological Services; 2003.

48. Chandler S, Charman T, Baird G, Simonoff E, Loucas T, Meldrum D, et al. Validation of the social communication questionnaire in a population cohort of children with autism spectrum disorders. Journal of the American Academy of Child & Adolescent Psychiatry. 2007;46(10):1324–32.

